# Targeting MYCN upregulates L1CAM tumor antigen in MYCN-dysregulated neuroblastoma to increase CAR T cell efficacy

**DOI:** 10.1101/2024.01.27.576592

**Authors:** Laura Grunewald, Lena Andersch, Konstantin Helmsauer, Silke Schwiebert, Anika Klaus, Anton G. Henssen, Teresa Straka, Marco Lodrini, Sebastian G. Wicha, Steffen Fuchs, Falk Hertwig, Frank Westermann, Alice Vitali, Carlotta Caramel, Gabriele Büchel, Martin Eilers, Kathy Astrahantseff, Angelika Eggert, Uta E. Höpken, Johannes H. Schulte, Thomas Blankenstein, Kathleen Anders, Annette Künkele

**Author notes:** <u>Corresponding author:</u> Annette Künkele, MD Charité - Universitätsmedizin Berlin Department of Pediatric Oncology and Hematology Augustenburger Platz 1 13353 Berlin Germany.

## Abstract

**Background:** Current treatment protocols have only limited success in pediatric patients with neuroblastomas harboring amplifications of the central oncogene, *MYCN*. Adoptive T cell therapy presents an innovative strategy to improve cure rates. However, L1CAM-targeting CAR T cells achieved only limited response against refractory/relapsed neuroblastoma in an ongoing phase I trial to date. Here, we investigate how oncogenic MYCN levels influence tumor cell response to CAR T cells, as one possible factor limiting success in trials.

**Methods:** High MYCN levels were induced in SK-N-AS cells harboring the normal diploid *MYCN* complement using a tetracycline-inducible system. The inducible MYCN cell model or *MYCN*-amplified neuroblastoma cell lines were cocultured with L1CAM-CAR T cells. CAR T cell effector function was assessed via activation marker expression (flow cytometry), cytokine release and tumor cytotoxicity (biophotonic signal assessment). The cell model was characterized using RNA sequencing, and our data compared to publicly available RNA and proteomic data sets from neuroblastomas. ChIP-sequencing data was used to determine transcriptional *L1CAM* regulation by MYCN using public data sets. Synergism between CAR T cells and the MLN8237 AURKA inhibitor, which indirectly inhibits MYCN activity, was assessed *in vitro* using the Bliss model and *in vivo* in an immunocompromised mouse model.

**Results:** Inducing high MYCN levels in the neuroblastoma cell model reduced L1CAM expression and, consequently, L1CAM-CAR T cell effector function (activation, cytokine release and cytotoxicity) *in vitro*. Primary neuroblastomas possessing high *MYCN* levels expressed lower levels of both the *L1CAM* transcript and L1CAM tumor antigen. Indirectly inhibiting MYCN via AURKA using MLN8237 treatment restored L1CAM expression on tumor cells *in vitro* and restored L1CAM-CAR T cell effector function. Combining MLN8237 and L1CAM-CAR T cell treatment synergistically increased neuroblastoma-directed killing in MYCN-overexpressing cells *in vitro* and *in vivo* concomitant with severe *in vivo* toxicity.

**Conclusion:** We shed new light on a primary resistance mechanism in MYCN-driven neuroblastoma against L1CAM-CAR T cells via target antigen downregulation. These data suggest that combining L1CAM-CAR T cell therapy with pharmacological MYCN inhibition may benefit patients with high-risk neuroblastomas harboring *MYCN* amplifications.

## Background

The International Neuroblastoma Risk Group (INRG) bases neuroblastoma risk stratification on different molecular markers, age group and genetic factors (1). *MYCN* amplification is the most common recurrent genetic aberration, occurring in ∼20% of neuroblastomas, in which it alone confers high risk and is associated with poor survival (2). High MYCN levels in neuroblastomas lacking *MYCN* amplifications also correlate with poor prognosis (3), and in mouse models, drive tumor development (4), linking oncogenic function to high MYCN levels.

Aberrant tumor suppressor or oncogene expression has only recently been demonstrated to alter a cancer’s ability to shape the host immune response to cancer (5, 6). High-risk *MYCN*-amplified tumors are immunologically “cold”, promoting a T cell-poor environment. Immunohistochemical quantification in archived neuroblastoma samples confirmed that *MYCN* amplification correlates with significantly lower CD4^+^ and CD8^+^ T cell infiltration (7). IFNG signaling and T cell-attracting chemokine (CXCL9, CXCL10) release are negatively regulated in primary neuroblastomas with oncogenic MYCN levels and *MYCN*-driven murine neuroblastic tumors, indicating oncogenic MYCN levels could hamper T cell infiltration and responsiveness (6).

Conventional therapies achieve only limited success against high-risk neuroblastoma, with <30% overall survival (8) that declines to <20% in patients with recurrent disease (9). An innovative therapeutic approach is chimeric antigen receptor (CAR) T cell therapy, which hijacks the immune system to direct T cell effector mechanisms against tumor cells. CARs are synthetic receptors usually constructed by linking a single-chain variable fragment (scFv) from a monoclonal antibody recognizing a tumor-specific protein on the cell surface to a transmembrane domain as well as no (1^st^ generation), one (2^nd^ generation) or more (3^rd^ generation) intracellular T cell costimulatory signaling modules (4-1BB or CD28) and the CD3ζ T cell signaling domain (10). Our group developed CAR T cells targeting the glycosylated CE7 epitope of L1CAM (11–13), which is specifically expressed on neuroblastoma cells (12). The ongoing phase I trial (NCT02311621) treats children with primary refractory or relapsed neuroblastoma with L1CAM-CAR T cells. Limited responses were observed in the first five enrolled patients, who all had *MYCN*-amplified disease (12).

Here we investigated how oncogenic *MYCN* levels influence L1CAM-CAR T cell effector function to better understand the unsatisfactory clinical outcomes achieved in the ongoing clinical trial (NCT02311621) for patients with primary refractory or relapsed neuroblastoma. We used neuroblastoma cell models allowing tight regulation of MYCN levels to create normal and oncogenic MYCN levels in different molecular cellular backgrounds. MYCN-mediated influence on tumor escape from CAR T cell therapy was preclinically analyzed in these models and in combination with indirect pharmacological MYCN inhibition as a potential treatment strategy for children with high-risk neuroblastomas harboring *MYCN* amplifications.

## Material and Methods

### Mice

Male and female NOD/SCID/γc^−/−^ (NSG) mice were group-housed according to institutional guidelines, compliant with national and EU regulations for animal use in research. Age- and sex-matched mice were subcutaneously injected (right flank) on day 0 with induced/uninduced 5×10^6^ SK-N-AS-TR-MYCN tumor cells in 50µl Matrigel™ (Corning)/50µl phosphate-buffered saline. Mice with induced SK-N-AS-TR-MYCN tumor cells received 0.2mg/ml doxycycline (a more stable form of tetracycline) in 5% sucrose-supplemented drinking water throughout the experiment. When tumors were palpable, mice were ranked by tumor size on the day of CAR T cell treatment, and treatment groups were randomized and contained mice with similar mean tumor sizes. Mice intravenously received 1×10^7^ untransduced or L1CAM-28/ζ CD3^+^ T cells in 100µl phosphate-buffered saline. MLN8237 (15mg/kg/mouse) was administered by oral gavage twice daily within a treatment regimen of 5 days on/2 days off for combination experiments. Mice were sacrificed when tumors reached a 1.500mm^3^ mean, according to *Landesamt für Gesundheit und Soziales (LAGeSo)* Berlin. Tumor material was processed using a previously described protocol (14).

### Cell lines and models

Neuroblastoma cell lines, SK-N-AS, SK-N-SH and IMR5/75, were cultured in RPMI 1640 (ThermoFisher Scientific) while SK-N-BE(2) and SK-N-DZ were maintained in Dulbeccós Modified Eagle Medium (ThermoFisher Scientific). The MYCN-inducible cell models were cultured in the same medium base as parental cell lines, but additionally supplemented with 100U/ml penicillin-streptomycin (Gibco), 5µg/ml blasticidin and, for SK-N-AS-MYCN, 500µg/ml G418-BC (Merck) or, for IMR5/75-shMYCN, 50µg/ml zeocin (ThermoFisher Scientific) as selection antibiotics (15). Cell models were induced by 2µg/ml tetracycline in full medium. Medium for neuroblastoma cell lines and models was supplemented with 10% heat-inactivated fetal calf serum (Sigma-Aldrich), and cultures were maintained at 37°C in 5% CO_2_. Neuroblastoma cell lines were transduced with a lentivirus encoding a GFP_firefly luciferase epHIV7 construct, producing a biophotonic light signal (for use in the cytotoxicity assay) and GFP expression. Cell lines were regularly checked for *Mycoplasma* contamination using the cell-based colometric HEK-Blue detection assay (Invivogen) and passaged maximally 20 times. Cell line identity was confirmed by STR fingerprinted (Eurofins, Luxemburg).

### CAR T cell generation

Magnetic-activated cell separation isolated CD8^+^ and CD3^+^ T cell populations [CD8^+^ T cell isolation kit as previously described (11, 16) or Pan T Cell Isolation Kit for CD3^+^ selection; Miltenyi Biotec] from human peripheral blood mononuclear cells from healthy donors (Charité ethics approval EA2/262/20). CD8^+^ L1CAM-CAR T cells (used for *in vitro* experiments) were generated and expanded as previously described (16) and supplied with 0.5ng/ml IL15 (Miltenyi Biotec) and 50U/ml IL2 (Novartis). CD3^+^ T cells (used for *in vivo* experiments) were supplied with 0.5ng/ml IL15 (Miltenyi Biotec) and 10ng/µl IL7 (Miltenyi Biotec) (17). CAR constructs were linked downstream to a T2A self-cleaving peptide and truncated EGFR for CAR T cell detection and cetuximab immunomagnetic positive selection (enrichment). Untransduced T cells were used as negative controls (for CAR T cell treatment) in experiments.

### Flow cytometric analyses

Cell surface expression of CD3, CD8A, CD4 (all BioLegend) and L1CAM (Miltenyi Biotec) was detected by fluorophore-conjugated monoclonal antibodies on a Fortessa X-20 (BD Biosciences) 4-laser flow cytometer. Truncated EGFR expression was detected using biotinylated cetuximab (Bristol-Myers Squibb) and a phycoerythrin (PE)-conjugated streptavidin antibody (BioLegend). T cell activation was assessed by fluorophore-conjugated monoclonal antibodies detecting CD137 (also known as TNFRSF9; BioLegend) and CD25 (also known as IL2RA; BioLegend). Dead cells were excluded from analyses using the LIVE/DEAD^TM^ Fixable Red Dead Cell Stain Kit (Life Technologies). GFP-expressing neuroblastoma cells were identified through the FITC channel. QuantiBRITE PE calibration beads (BD Biosciences) were used to determine L1CAM antigen density on neuroblastoma cells according to manufacturer’s instructions. Data was processed using FlowJo_V10 Software (Tree Star Inc.).

### Cytotoxicity assay

*In vitro* CAR T cell-mediated tumor cytotoxicity was quantified by luciferase-based reporter assay as previously described (18). For combinatorial treatments, CAR T cells were added to achieve indicated effector:target (E:T) ratios together with MLN8237 (Axon Medchem) inhibitor concentrations (1, 15, 25, 40, 60, 80, 100, 700 and 2,000nM) added from a 10mM stock solution (in DMSO) by the Tecan D300e Digital Dispenser (HP) for accurate volume delivery. Xenolight D-luciferin (PerkinElmer Inc.) was added (0.14mg/ml) after 72h treatment, and the biophotonic signal quantified (Promega GloMax Multi) after 3min. Tumor cell lysis mediated by combination treatment was determined using the formula, %specific lysis = (1- [RLU_sample_/RLU_max_])x100%, in relation to untreated tumor cells.

### Cytokine assays

IL2 and IFNG release from untransduced and CAR T cells was quantified in media conditioned for 24h by cocultures with neuroblastoma cell lines as previously described (18). Neuroblastoma cell lines were seeded for combination treatments at 5x10^5^ cells/well into 48-well plates with untransduced or L1CAM-CAR T cells (E:T ratio of 1:10) and 40nM of MLN8237.

### Quantitative Real-Time PCR (qRT-PCR)

Using the RNeasy Mini Kit (Qiagen), mRNA was isolated from cells and reverse transcribed into cDNA using the Transcriptor First Strand cDNA Synthesis Kit (Roche). Gene expression was quantified using FastStart Roche SybrGreen (Roche), the SteponePlus™ Real-Time PCR System (Applied Biosystems) and primer pairs (Eurofins) detecting RNA28S1 (fwd: TTGAAAATCCGGGGGAGAG, rev: ACATTGTTCCAACATGCCAG) and L1CAM (fwd: CATCCAGTAGATCCGGAG, rev: CTCAGAGGTTCCAGGGCATC). Delta CT was calculated relative to the *RNA28S1* housekeeping gene using StepOne Software v2.3 (Applied Biosystems)

### Droplet digital PCR

*MYCN* copy numbers were determined as previously described (19).

### Immunoblotting

Whole tumor cells were lysed 30 min on ice in 15mM HEPES, 150mM NaCl, 10mM EDTA, 2% Triton-X100 with Roche protease inhibitor and phosphatase inhibitor cocktails. The RotiQuant Bradford assay (Roth) determined protein concentrations and 10 or 20µg of protein were separated on 10% SDS-PAGE for western blotting. Proteins were detected using mouse monoclonal antibodies detecting MYCN (B8.4B; sc-53993, Santa Cruz Biotechnology), L1CAM (UJ127.11; ThermoFisher Scientific) or GAPDH (sc-32233, Santa Cruz Biotechnology) together with horseradish peroxidase-conjugated mouse IgG (diluted 1:5,000, Dianova). Proteins were visualized using the Fusion FX system and fold change was calculated using VisionCapt_v16.16d.

### RNA Sequencing

Total RNA sequencing libraries were generated as previously described (20). Briefly, RNA was extracted from tumor cells using Trizol™ (ThermoFisher Scientific), followed by enzymatic ribosomal RNA depletion, then transcribed, fragmented and hybridized using the TrueSeq Stranded mRNA kit (Illumina, San Diego, CA, USA). Libraries were sequenced on a HiSeq4000 sequencer (Illumina) with a paired-end read length of 2x150nt and a sequencing depth of 100 million reads by the Max Delbrueck Center for Molecular Medicine Sequencing Core (Berlin).

### Data analysis

Public microarray expression data (R2 genomics and visualization platform, http://r2.amc.nl/) from primary neuroblastoma cohorts (21) were re-analyzed to identify the relationship between *MYCN* and *L1CAM* expression. Chromatin immunoprecipitation sequencing (ChIP-seq) GSE80151 (22, 23) and GSE94782 (22, 23) datasets were downloaded from the Gene Expression Omnibus. Data quality was controlled (FASTQC 0.11.8) and adapters trimmed (BBMap 38.58) before aligning reads to the human hg19 consensus genome assembly (BWA-MEM 0.7.15, default parameters) and removing duplicate reads (Picard 2.20.4). ChIP-seq mappings were quality controlled with normalized & relative strand cross-correlation (NSC & RSC) (Phantompeakqualtools 1.2.1). Only data with NSC>1.05 and RSC>0.8 were further analyzed, following ENCODE recommendations (24). Reads were extended to 200bp (BigWig tracks in Deeptools 3.3.0) while masking blacklisted regions (ENCODE) and normalizing 10bp bins to counts/million before calling peaks (MACS2 2.1.2, default parameters). Combinatorial treatment (CAR T+MLN8237) synergy scores were calculated using R 1.2.5033 and the SynergyFinder package (25). E:T ratios (CAR T:tumor cells) were transformed into concentrations in MLN8237 IC50 range (1:1 E:T ratio = 1,000nM, 1:2 = 500nM, 1:5 = 200nM, 1:10 = 100nM) to analyze tumor cytotoxicity data (3 biological replicates). Data points for CAR T cells or inhibitor alone and in combination were used as default input to plot data and analyze synergism without bias [R using the Bliss model (26)] and visualize the drug combination dose-response landscape.

### Statistics

Significant differences in CAR T cell activation, cytokine release and tumor cytotoxicity (compared to untransduced T cells) were determined in paired and unpaired Student’s *t*-tests. Gene and protein expression data from neuroblastomas and L1CAM surface expression (*in vitro*) were compared using one-way ANOVA. Mouse cohorts treated with L1CAM-specific CAR T cells or untransduced T cells were compared using Kaplan–Meier survival analysis with log-rank statistics. Statistical analyses were conducted using GraphPad Prism 8 Software (GraphPad). Results were considered significant if p≤0.05.

## Results

### Oncogenic MYCN expression impairs L1CAM-CAR T cell effector function

Oncogenic *MYCN* amplification has been linked to a T cell-poor tumor microenvironment (6), providing a rationale to analyze how *MYCN* amplification influences CAR T cell efficacy. To study the impact of tightly regulated oncogenic *MYCN* levels, we used the L1CAM^+^ SK-N-AS neuroblastoma cell line (diploid *MYCN*) equipped with tetracycline-inducible MYCN expression (15). High MYCN levels were confirmed following induction with tetracycline and were comparable to amplified neuroblastoma cell lines harboring ∼100 *MYCN* copies per cell (**Figure S1A**). We assessed the relative ability of cocultured SK-N-AS-MYCN cells (±MYCN induction) to activate effector function in L1CAM-CAR T cells harboring either a 4-1BB (L1CAM-BB/ζ) or CD28 (L1CAM-28/ζ) costimulatory domain. Inducing oncogenic MYCN levels in the SK-N- AS-MYCN cells reduced activation (lower CD25 and CD137 expression, **Figure 1A**; gating strategy shown in **Figure S1B**) and effector function (IFNG and IL2 release, **Figure 1B**) in cocultured L1CAM-CAR T cells but did not trigger any cytokine release by untransduced T cells, confirming no activation in the negative control and antigen dependency of T cell activation. Transduced CAR T cells were enriched by cetuximab immunomagnetic positive selection (binds the truncated EGFR tag) to 97.5% (L1CAM-BB/ζ) and 98.6% (L1CAM-28/ζ, **Figure S1C**). Viable tumor cells (±MYCN induction) were quantified by reporter assay following CAR T cell exposure to assess how MYCN influences CAR T cell-mediated cytotoxicity. L1CAM-CAR T cell-dependent tumor cytotoxicity was reduced by high-level MYCN induction (**Figure 1C**), regardless of the costimulatory domain used in the CAR construct (4-1BB: 25.8%±1.2% vs. 36.2%±3.4%; CD28: 60.9%±6.2% vs. 81.7%±5.6%).

**Fig. 1:**
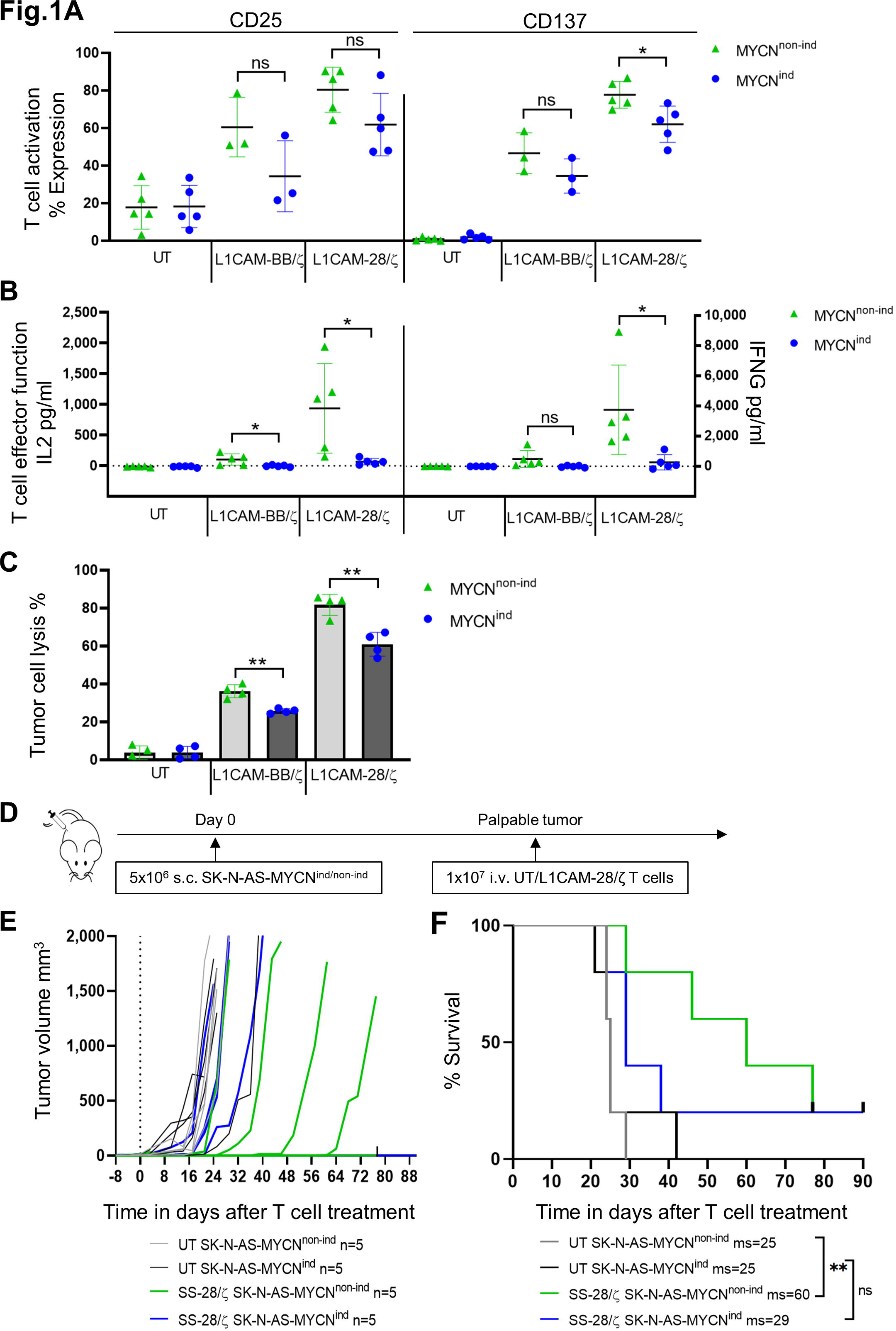
L1CAM-CAR T cell display impaired effector function when stimulated with MYCN^ind^ neuroblastoma cell model *in vitro* and *in vivo*. A. CD25 and CD137 surface molecule expression on viable CD8^+^ L1CAM-CAR and untransduced (UT) T cells after 24 h coculture with SK-N-AS-MYCN^non-ind/ind^ tumor cells (effector to target ratio (E:T) of 1:5, n=3 L1CAM-BB/ζ, n=5 L1CAM-28/ζ) measured by flow cytometry. **B.** IL2 and IFNG cytokine release by CAR T cells cocultured with SK-N-AS-MYCN^non-ind/ind^ cells for 24 h was analyzed using ELISA (E:T 1:5; n=5 biological replicates with each in technical triplicates). **C.** SK-N-AS-MYCN^non-ind^ tumor cells were stably transduced with GFP_fflluc and cocultured with L1CAM-CAR T cells (E:T 1:5). Tumor cell lysis was determined by a luciferase-based reporter assay relative to an untreated coculture after 48 h (n=4 biological replicates with each in technical triplicates). **D.** Scheme of experimental set-up. **E.** Tumor growth curves of NSG mice harboring either SK-N-AS-MYCN^non-ind^ or -MYCN^ind^ tumors treated with untransduced (UT) and L1CAM-28/ζ CAR T cells (all groups n=5). Each line represents changes in tumor volume of an individual mouse over time of the experiment. Zero (“0”) indicates start of treatment. **F.** Kaplan-Meier survival curve of NSG mice harboring either SK-N-AS-MYCN^non-ind^ or -MYCN^ind^ tumors treated with untransduced (UT) and L1CAM-28/ζ CAR T cells (all groups n=5; log-rank Mantel-Cox test) Zero (“0”) indicates start of treatment. mean ± SD, students T-test, ns = not significant, *, p≤0.05; **, p≤0.01.; UT=untransduced

Untransduced T cell controls killed <4% of cocultured tumor cells. To extend our analysis to the *in vivo* situation, subcutaneous tumors derived from inducible SK-N-AS-MYCN tumor cells were initiated in NSG mice (followed by ±MYCN induction, **Figure S1D**), then injected with L1CAM-28/ζ CAR T cells, which responded more strongly in *in vitro* testing (**Figure 1D**). MYCN induction alone did not alter tumor growth (**Figure 1E**), indicating that MYCN expression level did not influence growth kinetics in our model. CAR T cell injection had no effect on growth of tumors with oncogenic MYCN levels, but delayed growth in 3 of 5 tumors with normal MYCN levels. Median survival (MS) of mice harboring uninduced tumors and challenged with CAR T cells was significantly longer than mice challenged with untransduced T cells (MS: 60 vs 25 days; p=0.0048). Survival was not significantly enhanced (than untransduced T cell controls) after CAR T cell challenge in mice with tumors expressing oncogenic MYCN levels (MS: 29 vs 25 days; **Figure 1F**). Results using this neuroblastoma model with tightly regulable MYCN levels support that oncogenic MYCN levels in neuroblastoma reduces L1CAM-CAR T cell effector functions.

### Inducing high-level MYCN expression diminishes L1CAM surface expression on neuroblastoma cells

We next sought to unravel the mechanism underlying MYCN-mediated impairment of CAR T cell efficacy. Global transcription profiles were analyzed in the SK-N-AS-MYCN cell model before and 48h after inducing high MYCN levels. Induction increased *MYCN* transcript levels by >6-fold. Oncogenic MYCN levels upregulated (by almost 2-fold) MAX interactor 1, dimerization protein (MXI1), which competes with MYCN for MAX binding to mediate transcriptional repression (27), and downregulated a number of genes expressed on the cell surface (**Figure 2A**). These including L1 family members, *NFASC*, *CD177* and *CDH6*, as well as our CAR T cell target antigen, *L1CAM*, which was among the most downregulated genes (>2-fold). Flow cytometric evaluation of L1CAM expression on the SK-N-AS-MYCN cell surface 3 days after inducing high MYCN levels showed L1CAM expression to be diminished by 1.7-fold (p=0.0098; **Figure S2A**), and bead-based quantification detected a 10-fold reduction in L1CAM molecules on the SK-N-AS-MYCN cell surface (**Figure 2B**). We explored whether the negative correlation between oncogenic MYCN expression and L1CAM surface expression was recapitulated in other MYCN-regulable neuroblastoma cell models. SK-N-SH-MYCN cells, harboring the same tetracycline MYCN-inducible system (15), displayed significantly fewer (1.8-fold) L1CAM molecules per cell after inducing high MYCN levels (**Figure 2B**). *MYCN* knockdown using tetracycline-inducible shRNA in IMR5/75-shMYCN cells, which harbor a high-level *MYCN* amplification, increased L1CAM cell surface molecules by 1.4-fold (**Figure 2B**). Changes in L1CAM expression in all 3 inducible models were confirmed on the transcript level (**Figure 2C**). We assessed whether the influence of oncogenic MYCN levels was sustained on target antigen expression over time in the SK-N-AS-MYCN model. L1CAM cell surface expression (assessed before and 3, 7 and 14 days after inducing MYCN) was further reduced to 3.6-fold by day 14 of sustained high MYCN levels (**Figure S2B**), and reduced effector cytokine release by L1CAM-CAR T cells correlated with reduced L1CAM surface expression on neuroblastoma cells (**Figure S2C**). Our results from MYCN-regulable cell models confirm that oncogenic MYCN levels reduce L1CAM transcription and expression on the neuroblastoma cell surface.

**Fig. 2:**
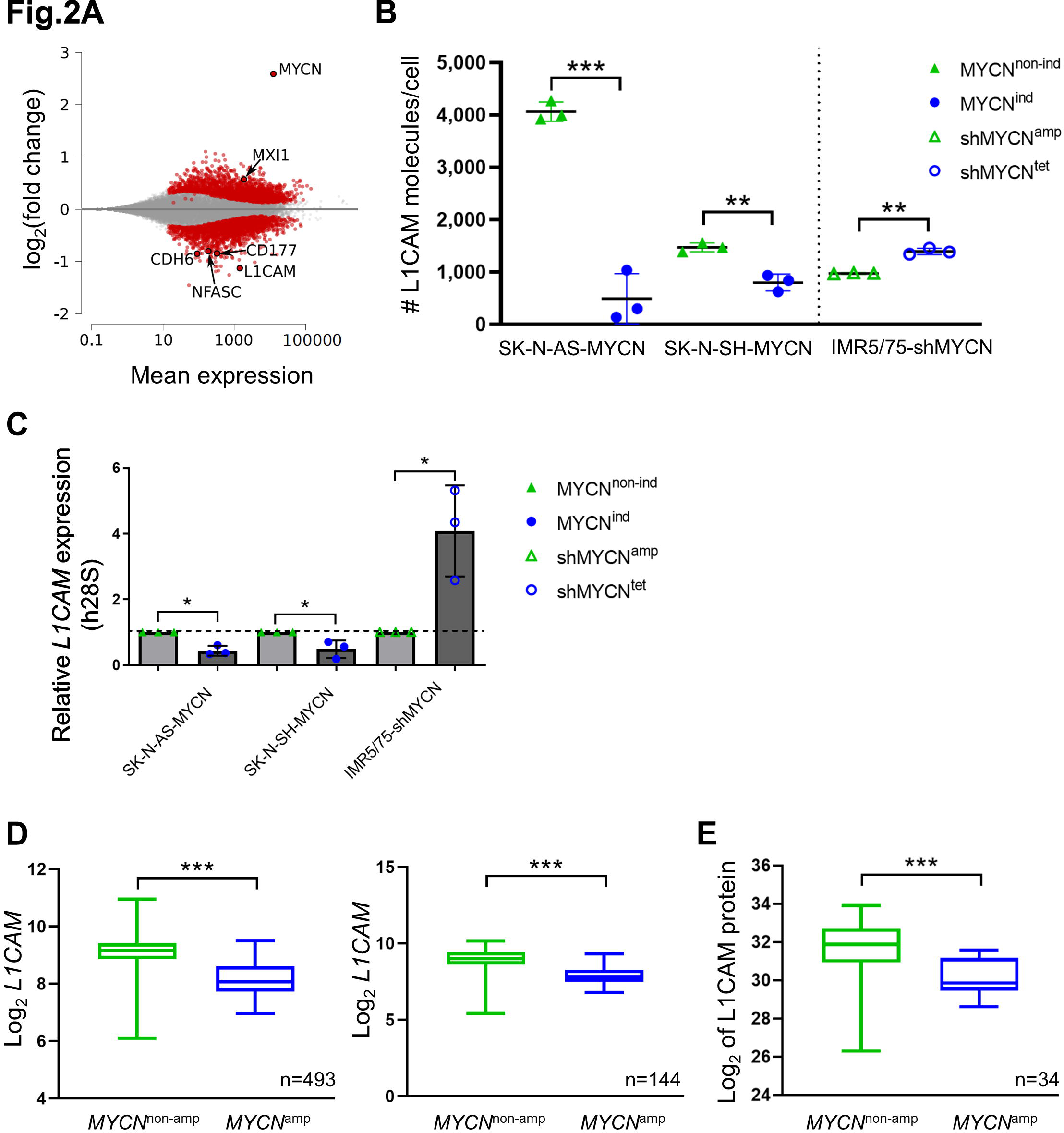
M**Y**CN **overexpression correlates with reduced L1CAM expression on neuroblastoma cells and in patient cohorts. A.** MA-Plot of RNA-sequencing data using DESeq2 of fold change (log_2_) of MYCN induction in SK-N-AS-MYCN^non-ind^ versus MYCN^ind^ cells after 48 h of tetracycline treatment. Red dots represent up- or downregulated genes upon MYCN induction (n=3 biological replicates, M (log ratio), A (mean average)). **B.** L1CAM cell surface expression quantified on SK-N-AS-MYCN^non-ind^, - MYCN^ind^, SK-N-SH-MYCN^non-ind^, -MYCN^ind^, IMR5/75-shMYCN^amp^ and -MYCN^tet^ tumor cells. Data shows L1CAM molecules per cell (n=3 biological replicates). **C.** Relative expression *L1CAM* mRNA levels in SK-N-AS-MYCN^non-ind/ind^, SK-N-SH-MYCN^non-ind/ind^ and IMR5/75-shMYCN^amp/tet^ tumor cells relative to housekeeper h28S. **D.** Gene-expression data from two patient cohort represents log_2_ fold change of *L1CAM* expression in a cohort of 498 neuroblastoma patients, without (*MYCN*^non-amp^ n=401) and with *MYCN* amplification (*MYCN*^amp^ n=92, n=5 no *MYCN* status available, not included, ANOVA p=4.53e-30) and in a cohort of 144 neuroblastoma patients, without (*MYCN*^non-amp^ n=104) and with *MYCN* amplification (*MYCN*^amp^ n=40, ANOVA p=2.63e-12) (21). **E.** Patient data from Hartlieb et al. represents log_2_ change of L1CAM protein expression in a cohort of 34 neuroblastoma patients, without (*MYCN*^non-amp^ n=22) and with *MYCN* amplification (*MYCN*^amp^ n=12, ANOVA p=7.31e-3) (21).

To validate the clinical relevance of our findings from MYCN-regulable neuroblastoma cell models, we re-analyzed datasets for *L1CAM* expression in primary neuroblastoma samples from 2 independent patient cohorts, each containing tumors harboring or lacking *MYCN* amplifications. Microarray-based gene expression data profiles from 493 neuroblastomas (28) and RNA sequencing data from 144 neuroblastomas (21) confirmed that *L1CAM* expression was 2-fold lower in neuroblastomas harboring *MYCN* amplifications than tumors lacking *MYCN* amplifications in either cohort (**Figure 2D-E**). L1CAM protein abundance was also significantly downregulated (p=0.0073) in *MYCN*-amplified primary neuroblastomas in publicly available mass spectrometry sequencing data (21) (**Figure 2E**, n=34). We also reanalyzed ChIP sequencing data from the *MYCN*-amplified neuroblastoma cell lines, SK-N-BE(2)-C, Kelly and NGP (22, 23), to assess whether MYCN may directly regulate *L1CAM* expression. MYCN peaks were identified within the *L1CAM* gene body, suggesting that it may be involved in its regulation (**Figure S3**). In two of three cell lines, we identified MYCN binding at the promotor, marked by H3K4me3. MYCN peaks also colocalized with an accessible chromatin region in the first intron, as indicated by ATAC-seq, which was also marked by H3K27 acetylation (**Figure S3**). This indicates that MYCN binds to a putative intronic enhancer in *L1CAM*, and may be involved in L1CAM regulation. We show that oncogenic MYCN levels achieved either through *MYCN* amplification or enhanced expression diminished L1CAM target protein on the neuroblastoma cell surface. This target reduction may be the mechanism behind resistance to L1CAM-CAR T cell therapy.

### Exogenous L1CAM surface expression restores *in vitro* CAR T cell effector activity

We explored whether restoring L1CAM expression on neuroblastoma cells is sufficient to rescue MYCN-mediated attenuation of CAR T cell function. Constitutive L1CAM surface expression was achieved in the SK-N-AS-MYCN model by transducing a lentiviral vector encoding the *L1CAM* transgene. High L1CAM expression, independent of MYCN level, was flow cytometrically confirmed in L1CAM-SK-N-AS-MYCN cells (**Figure 3A**). L1CAM-CAR T cells were similarly activated (**Figure 3B**), released similar levels of effector cytokines (**Figure 3C**) and mediated cytotoxicity (**Figure 3D**) when cocultured with L1CAM-SK-N-AS-MYCN cells regardless of MYCN induction. CAR constructs using either costimulatory domain achieved similar activities. These data support that enforcing L1CAM expression is sufficient to rescue CAR T cell efficacy against the tumor cells even in the presence of the escape mechanism driven by oncogenic MYCN levels.

**Fig. 3:**
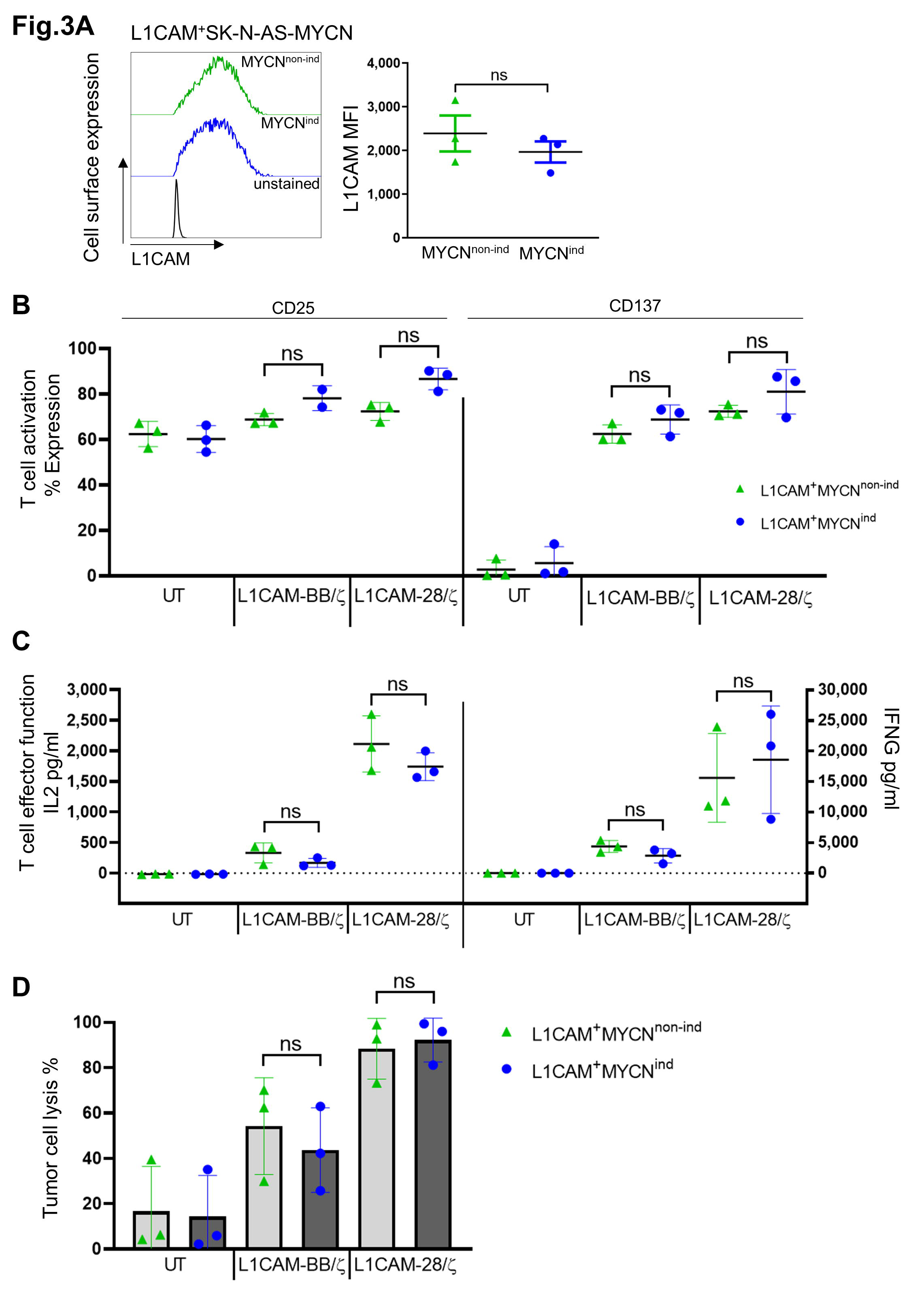
S**t**able **L1CAM expression restores CAR T cell effector function against tumor cells with MYCN overexpression. A.** L1CAM surface expression was analyzed on SK-N-AS-MYCN tumor cells engineered to express L1CAM under a strong constitutively active promoter EF1A (L1CAM^+^SK-N-AS- MYCN). Cells were either cultured without (MYCN^non-ind^) or with 2µg/ml tetracycline (MYCN^ind^). Exemplary flow cytometry staggered histogram plots per condition and dot blots representing L1CAM MFI are shown (n=3 biological replicates). **B.** Activation marker expression levels on L1CAM-CAR T cells after 24 h coculture with L1CAM^+^SK-N-AS-MYCN^non-ind^ or -MYCN^ind^ cells (E:T 1:5; n=3 biological replicates). **C.** Quantification of cytokine release after a 24 h coculture of L1CAM^+^SK-N-AS- MYCN^non-ind^ -MYCN^ind^ tumor cells with L1CAM-specific CAR T cells (E:T 1:5; n=3 biological replicates in technical triplicates). **D.** L1CAM^+^SK-N-AS-MYCN^non-ind/ind^ tumor cell lysis assay after coculture with L1CAM-specific CAR T cell was determined by a luciferase-based reporter assay relative to an untreated coculture after 48 h (E:T 1:5, n=3 biological replicates in technical triplicates). Mean ± SD, students T- test, ns = not significant.

### MLN8237 treatment enhances neuroblastoma L1CAM expression to boost L1CAM-CAR T cell efficacy

We investigated whether pharmacologically inhibiting MYCN activity could produce the same effect as enforcing L1CAM expression, and restore L1CAM-CAR T cell neuroblastoma cytotoxicity. The alpha-helical structure of the MYCN transcription factor has prevented development of direct MYCN-targeting agents so far. MLN8237 is a small molecule that indirectly inhibits MYCN by targeting the aurora A kinase (AURKA) to drive MYCN degradation (**Figure 4A**) (29). MLN8237 treatment (80, 800nM) dose-dependently reduced MYCN levels in SK-N-AS-MYCN cells (±MYCN induction, **Figure S4A**). MYCN reduction in SK-N-AS-MYCN cells (±MYCN induction) treated with the lower MLN8237 dose enhanced L1CAM surface expression by 1.2-fold, while treatment with 800nM MLN8237 enhanced L1CAM surface expression by 1.4-fold (low MYCN levels) and 1.6-fold (high MYCN levels, **Figure 4B**). Since oncogenic MYCN levels appeared to impact CAR T cells utilizing CD28 costimulation slightly more strongly (**Figure 1**) and L1CAM-28/ζ CAR T cells were previously shown to be more effective against solid tumors (30), further testing was performed only with this CAR T cell product. MLN8237 treatment did not alter IL2 or IFNG cytokine release from CAR T cells exposed to SK-N-AS-MYCN (±MYCN induction) cells (**Figure S4B**). The combined effect of MLN8237 treatment (8 doses, range: 15-2,000nM) with L1CAM-CAR T cells (5 E:T ratios, range: 1:1-1:20) was tested on cocultured SK-N-AS-MYCN cells (±MYCN induction), using the Bliss independence model (26) to determine additive, synergistic or antagonistic effects. We have chosen the Bliss model as per definition both drugs used here, act independently against tumor cells and there is no drug-drug interaction (31). The combination of both low-dose treatments enhanced tumor cytotoxicity against tumor cells with oncogenic MYCN levels. (**Figures 4C-D**). To illustrate effects on SK-N-AS-MYCN cells with oncogenic MYCN levels more clearly, we depict each therapy alone and in combination at the peak synergy score (E:T = 1:10, 40nM MLN8237, **Figure 4E**), based on absolute inhibitory values of individual concentrations used in combinational treatment (**Figure S4C-D**). At the peak synergy score, combining treatments significantly increased tumor cytotoxicity (74.7±6.1%), while single treatments with CAR T cells (29.7±11.1%, p=0.0037) or MLN8237 (46.4±11.0%, p=0.018) achieved only limited cytotoxicity. Combining MLN8237 treatment with CAR T cells did not provide a significant benefit against SK-N-AS-MYCN cells with normal MYCN levels. Pharmacologically inhibiting MYCN activity works in concert with L1CAM-CAR T cell-directed cytotoxicity against SK-N-AS-MYCN cells with oncogenic MYCN levels, resulting in significantly enhanced tumor cell lysis *in vitro*.

**Fig. 4:**
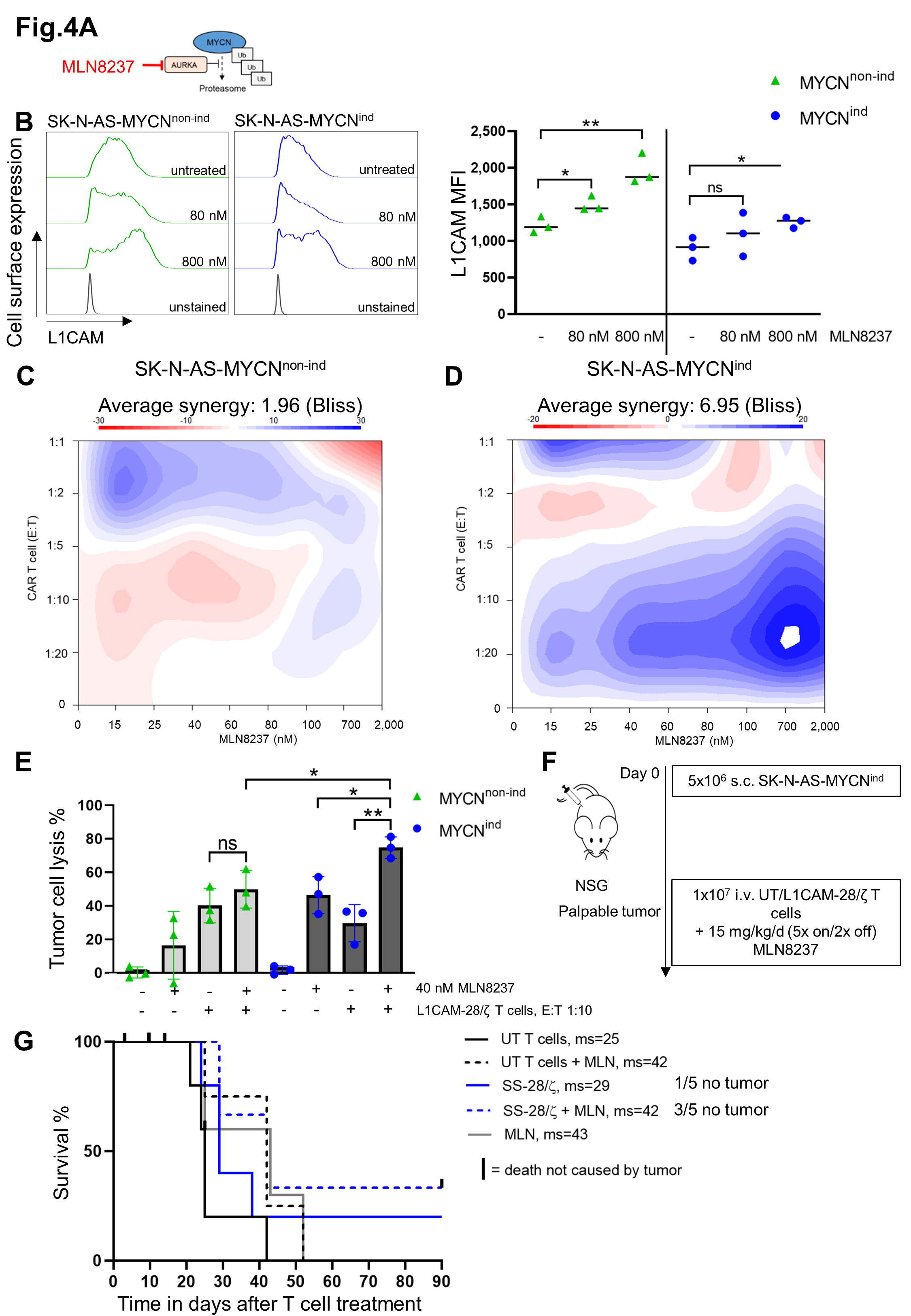
Indirect MYCN inhibition by MLN8237 increases L1CAM expression and improves L1CAM-CAR T cell effector function against MYCN^ind^ neuroblastoma cells *in vitro* and *in vivo*. A. Schematic overview of the mechanism of action of Aurora A kinase inhibitor MLN8237. **B.** L1CAM surface expression of SK-N-AS-MYCN^non-ind^ and -MYCN^ind^ tumor cells after treatment with 80 and 800nM MLN8237 for 72 h measured by flow cytometry. Dot blots represent MFI of L1CAM (n=3 biological replicates). **C.** and **D.** Bliss synergism model showing the heatmap indicating antagonism (red) and synergism in blue for SK-N-AS-MYCN^non-ind/ind^ tumor cells that were treated with L1CAM-CAR T cells and MLN8237 in different concentrations (n=3 in biological replicates). **E.** Cytolytic activity of L1CAM-28/ζ CAR T cells against SK-N-AS-MYCN^non-ind^ –MYCN^non-ind^ NB tumor cells (E:T 1:10) with or without 40nM MLN8237 (n=3 biological replicates in technical triplicates). **F.** Scheme of experimental set-up. **G.** Kaplan-Meier curve of NSG mice harboring SK-N-AS-MYCN^ind^ tumors treated with untransduced (UT) and L1CAM-28/ζ CAR T cells alone or in combination with MLN8237 (all groups n=5). Zero (“0”) indicates start of treatment. mean ± SD, students T-test, *, p≤0.05; **, p≤0.01.

We tested the combined L1CAM-CAR T cell and MLN8237 treatment against xenograft SK-N- AS-MYCN tumors with maintained oncogenic MYCN levels (doxycycline in drinking water) in our immunodeficient NSG mouse model. Once palpable tumors were detected, T cells (L1CAM- 28/ζ CAR or untransduced control T cells) were intravenously injected once into mice that either received MLN8237 or not by oral gavage twice daily in a cotreatment course of up to 90 days (**Figure 4F**). Inhibiting MYCN activity alone delayed tumor growth in 3 of 5 mice and improved median survival (MS) to 43 days, while L1CAM-CAR T cells alone did not delay tumor growth or improve survival compared to control mice treated with untransduced T cells (MS=29 days and 25 days, respectively; **Figure 4G**; **Figure S4E**). Combining L1CAM-CAR T cells with MLN8237 eradicated tumors in 3 of 5 mice and improved MS to 42 days. However, MLN8237 treatment caused severe toxicity in mice (discomfort not caused by tumor growth), resulting in the need to sacrifice 4 mice (1 treated with untransduced T cells + MLN8237 3 days after T cell injection, 2 treated with L1CAM-CAR T cells + MLN8237 14 days after T cell injection; 1 treated with MLN8237 after 25 days of treatment) and preventing statistical analysis of the combination effect on survival. This experiment revealed that although MLN8237 causes severe toxicity *in vivo*, overall survival appeared to improve by combining L1CAM-CAR T cells with inhibition of MYCN activity.

*MYCN* amplification may create a different cell background environment than raising MYCN expression to oncogenic levels alone in the MYCN-inducible model. We extended testing of combined treatment to 3 neuroblastoma cell lines harboring different *MYCN* copy numbers, IMR5/75 (112 copies), SK-N-DZ (130 copies) and SK-N-BE(2) with 487 copies **Figure S1A**), all considered *MYCN*-amplified. MLN8237 treatment significantly increased flow cytometrically detected L1CAM surface expression in IMR5/75 (2.0-fold) and SK-N-DZ (1.9-fold) cells, but not in SK-N-BE(2) cells (**Figure 5A**). L1CAM-CAR T cell-directed cytotoxicity was improved by MLN8237 cotreatment in IMR5/75 and SK-N-DZ cells, but not SK-N-BE(2) tumor cells, as would be expected by their lack of L1CAM target enhancement by MLN8237 (**Figure 5B**). MLN8237 does not appear to inhibit MYCN activity well in SK-N-BE(2), suggesting there may be some variability in the efficacy of co-treatment depending on the drug selected. Collectively, our findings demonstrate that combining inhibition of MYCN activity with L1CAM-CAR T cell therapy could increase the efficacy of L1CAM-CAR T cell therapy for patients with *MYCN*- amplified neuroblastoma by counteracting the MYCN-directed tumor escape mechanism that downregulates L1CAM target expression on the tumor.

**Fig. 5:**
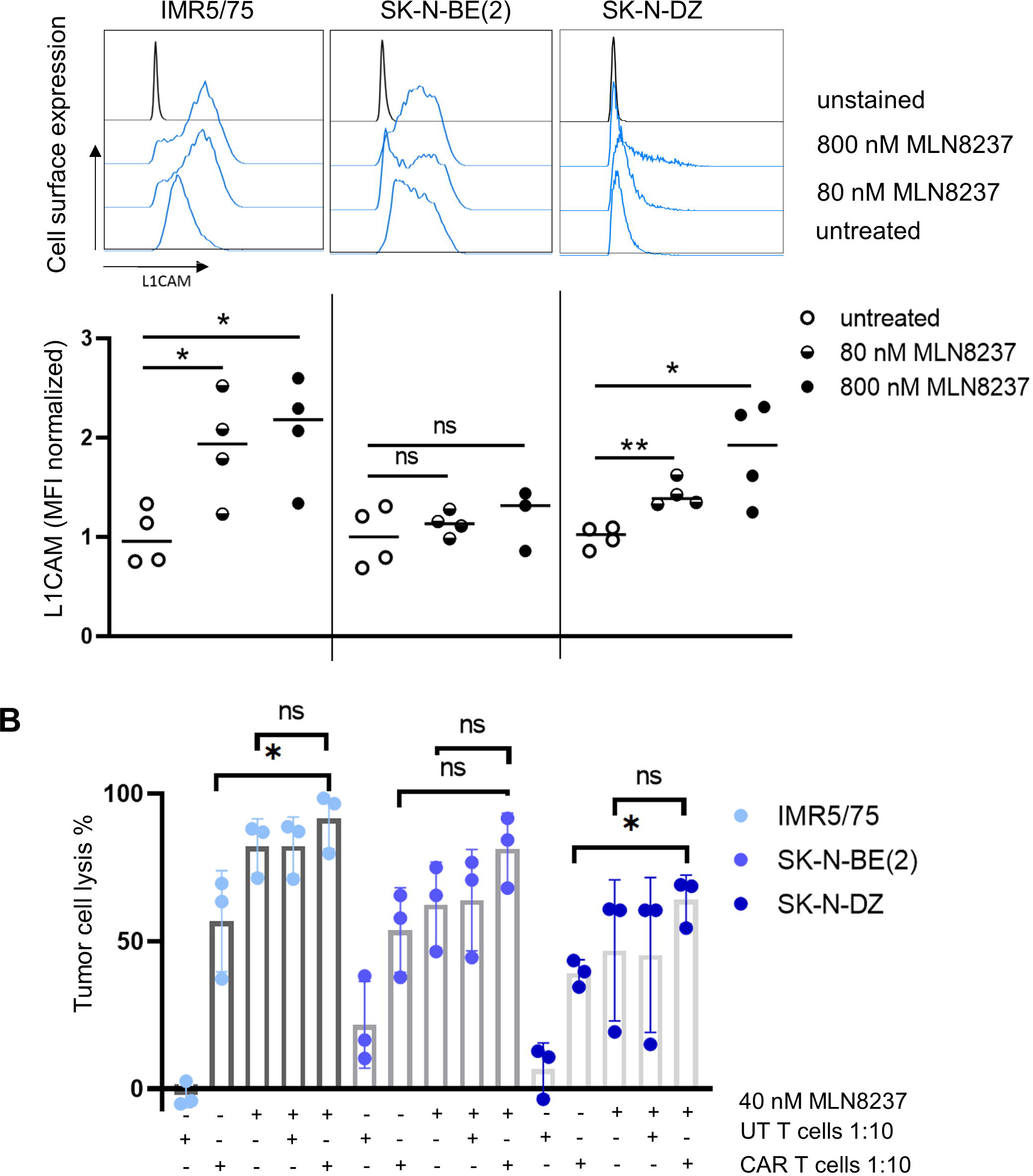
MLN8237 enhances L1CAM surface expression on *MYCN*-amplified tumor cell lines *in vitro*. **A.** Flow cytometry analysis representing L1CAM expression of IMR5/75, SK-N-BE(2) and SK-N-DZ cell lines treated with 80 and 800nM MLN8237 for 72 h. Dot blots represent normalized MFI of L1CAM of IMR5/75, SK-N-BE(2) and SK-N-DZ n=4. **B.** Combinational therapy of L1CAM-specific CAR T cells (L1CAM-28/ζ; E:T 1:10) and 40nM MLN8237. Biophotonic signal of IMR5/75, SK-N-BE(2) and SK-N- DZ cells was measured to analyze cytotoxic potential of therapies alone and in combination (n=3 biological replicates in technical replicates). mean ± SD, students T-test, ns= not statistically significant, *, p≤0.05.

## Discussion

Here we show that oncogenic MYCN levels in neuroblastoma impair L1CAM-CAR T cell efficacy by downregulating L1CAM target antigen expression on neuroblastoma cells. Combining the indirect MYCN inhibitor, MLN8237, with CAR T cells enhanced L1CAM-CAR T cell-directed cytotoxicity *in vitro* in neuroblastoma cells expressing oncogenic MYCN levels caused by induced upregulation in the diploid *MYCN* background or *MYCN* amplifications. Combined inhibition of MYCN activity and L1CAM-CAR T cell treatment also delayed neuroblastoma outgrowth in mice, in a background of tumor-unrelated toxicity to the MLN8237 inhibitor.

The presence of tumor infiltrating lymphocytes in many tumor entities positively correlates with improved clinical outcome [reviewed in (32), (7, 33)]. A T cell-poor microenvironment, with reduced IFNG signaling and chemokine activity (CXCL9 and CXCL10), characterizes oncogene-driven, *MYCN*-amplified neuroblastoma (6). This tumor environment is expected in the first five patients treated in the ongoing phase I clinical trial investigating L1CAM-CAR T cells, since *MYCN* amplifications were documented in diagnostic samples from all five patients (12), driving our aim to investigate how oncogenic MYCN levels impact CAR T cell efficacy. Here we demonstrate that oncogenic MYCN levels in neuroblastoma cells impair activation (reduced CD25 and CD137 molecules on T cells) and cytotoxic potential of L1CAM-CAR T cells to reduce effector function. Particularly IFNG effector cytokine release by L1CAM-CAR T cells declined severely with high MYCN levels, adding an immunosuppressive function to *MYCN* oncogene-driven tumor cells already previously shown to have a poor IFNG pathway activity by Layer *et al*. (6). Here, raising MYCN to oncogenic levels caused faster outgrowth in xenotransplanted tumors treated with L1CAM-CAR T cells and reduced mouse survival. Since *MYCN*-amplified tumors are known to harbor myeloid-derived suppressor cells and tumor-associated macrophages, which are immunosuppressive (34, 35), the tumor microenvironment could be contributing to the negative effect exerted on L1CAM-CAR T cells via oncogenic MYCN levels in the tumor cells. This cannot be tested in an immunocompromised NSG mouse model and is, therefore, a limitation in our study. We demonstrated that oncogenic MYCN drives L1CAM target antigen reduction and identified an inverse correlation of L1CAM surface expression and MYCN overexpression or *MYCN* amplification in neuroblastoma cell lines and patient data. This was in contrast to data from Rached *et al*. who reported that *L1CAM* knockdown reduced MYCN expression in *MYCN*-amplified IMR-32 neuroblastoma cells, with reductions in proliferation, migration and tumor sphere formation (36). CAR T cell effector function strongly depends on abundance of target antigen and is impaired when antigen expression levels decline below a certain threshold (37–39). Watanabe *et al*. demonstrated that CD20-CAR T cells effectively lysed tumor cells expressing ∼200 CD20 molecules/cell but required ∼5,000 CD20 molecules/cell to trigger effector cytokine production and T cell proliferation (40). Our results confirm that antigen density is pivotal for optimal L1CAM-CAR T cell efficacy, since reducing L1CAM molecules/cell from ∼4,000 to ∼500 by inducing oncogenic MYCN levels also diminished cytokine release and tumor cytotoxicity. This finding shows, to our knowledge, the first evidence that MYCN contributes to tumor escape from L1CAM-specific CAR T cells by downregulating the target antigen decreasing responsiveness or even causing primary resistance to L1CAM-CAR T cell therapy.

Our analyses showed that targeting MYCN with MLN8237 upregulates L1CAM surface expression on neuroblastoma cell lines dependent on MYCN copy numbers, subsequently improving L1CAM-CAR T cell efficacy. We also demonstrate how the Bliss model, only applied to data from drug combinations to date, can be modified to calculate synergism between CAR T cells and pharmacological inhibitors. The combination therapy delayed tumor outgrowth and seemed to improve overall survival of mice harboring tumors with high-level MYCN. However, survival could not be statistically analyzed because too many mice needed to be removed from the experiment due to toxicity. Discomfort of mice was only seen in MLN8237- treated animals (single treatment or in combination with untransduced T cells or CAR T cells), suggesting severe side effects of this drug *in vivo*. Our observation is in line with results by Mossé et al., who detected high toxicities, including myelosuppression, mucositis, neutropenia and depression among others in a recent phase II trial, where patients with recurrent/refractory solid tumors or leukemia received MLN8237 as a single agent (41). These toxicities have not been observed in the first preclinical evidence in mice, where MLN8237 produced a complete response against pediatric tumors and acute lymphoblastic leukemia independent of *MYCN* status (42). Alternative drug combinations with L1CAM-CAR T cells could be used, like indirect MYC family inhibitors, I-BET726 and JQ1, that show reduced immunogenicity profiles in neuroblastoma cells harboring *MYCN* amplifications, suggesting other MYC family inhibitors could have the same effect as MLN8237 but with lower toxicity profiles (43). These indirect inhibitors of MYCN, as well as next-generation AURKA inhibitors (e.g. LY3295668) are already used in clinical trials (NCT03936465, NCT04106219) making them available for future combination testing with L1CAM-CAR T cells in clinical trials.

Here we demonstrate that oncogenic MYCN levels impair L1CAM-CAR T cell effector function by reducing target molecules expressed on the neuroblastoma cells, providing a route to tumor immune escape. We provide preclinical evidence that pharmacologically inhibiting MYCN function restores L1CAM target expression on neuroblastoma cells. These findings offer the rationale for a future clinical trial to test the combination of MYCN-targeting drugs with L1CAM-CAR T cells in children with *MYCN*-amplified neuroblastoma.

## Declarations

### Ethics approval

Ethics approval for generating CAR T cells using T cells from healthy donors was obtained (Charité ethics committee approval EA2/262/20).

### Consent of publication

Not applicable.

### Availability of data and material

Data and materials will be provided by the corresponding author upon reasonable request.

### Competing interests

The authors declare no conflicts of interest.

### Funding

LG is participant in the “Berlin School of Integrative Oncology” and supported by the Wilhelm Sander Stiftung (#2021.115.1). AKü is a participant in the BIH – Charité Advanced Clinician Scientist Pilot Program. SF is a participant in the BIH-Charité Clinician-Scientist Program. All these programs are cofunded by the *Charité – Universitätsmedizin Berlin* and Berlin Institute of Health. This work was supported by the Else Kröner-Fresenius Foundation (#2017_A51 grant to AKü), *Deutsche Krebshilfe* (#701133871 grant to AKü, AGH and ME; #70112951 grant to AKü, ME, JHS, AE) and *Deutsche Forschungsgemeinschaft* (SFB-TRR 338/1 2021 –452881907 to AKü and ME; fellowship grant no. 439441203 to SF). The funders had no role in the study design, data collection and analysis, decision to publish or preparation of the manuscript.

### Authors’ contributions

LG designed and performed experiments, interpreted data and wrote manuscript. LA, SS, AKl, FZ, ML, TS, SF, AV and CC performed experiments. KH and AGH analyzed ChIP seq data. SGW helped with investigation of synergy scores. FW provided the IMR5/75-TR-shMYCN cell line. AGH, FH, GB, ME, AE, UEH, JHS and TB co-conceived study and interpreted data. AKü, JHS and AE acquired funding. AKü conceived and supervised study, KAn co-supervised study. AKü, KAn and KAs revised manuscript. All authors revised and approved the last version of the manuscript.

## Acknowledgments

We thank Michael C. Jensen for providing the L1CAM-CAR T cell constructs and Hans-Dieter Volk for providing the NSG mice.

## List of abbreviations

CAR: Chimeric antigen receptor
ChIP: Chromatin immunoprecipitation
ELISA: Enzyme-linked immunosorbent assay
E:T: Effector:target
INRG: International neuroblastoma risk grouping
MS: Median survival
NSC: Normalized strand cross-correlation
NSG: NOD/SCID/γc^-/-^
RSC: Relative strand cross-correlation

## Notes

### Competing Interest Statement

The authors have declared no competing interest.

## References

1. Cohn SL, Pearson AD, London WB, Monclair T, Ambros PF, Brodeur GM, et al. The International Neuroblastoma Risk Group (INRG) classification system: an INRG Task Force report. J Clin Oncol. 2009;27(2):289–97.

2. Westermann F, Muth D, Benner A, Bauer T, Henrich K-O, Oberthuer A, et al. Distinct transcriptional MYCN/c-MYC activities are associated with spontaneous regression or malignant progression in neuroblastomas. Genome Biology. 2008;9(10):R150.

3. Alaminos M, Gerald WL, Cheung NK. Prognostic value of MYCN and ID2 overexpression in neuroblastoma. Pediatr Blood Cancer. 2005;45(7):909–15.

4. Weiss WA, Aldape K, Mohapatra G, Feuerstein BG, Bishop JM. Targeted expression of MYCN causes neuroblastoma in transgenic mice. EMBO J. 1997;16(11):2985–95.

5. Dorand RD, Nthale J, Myers JT, Barkauskas DS, Avril S, Chirieleison SM, et al. Cdk5 disruption attenuates tumor PD-L1 expression and promotes antitumor immunity. Science. 2016;353(6297):399- 403.

6. Layer JP, Kronmuller MT, Quast T, van den Boorn-Konijnenberg D, Effern M, Hinze D, et al. Amplification of N-Myc is associated with a T-cell-poor microenvironment in metastatic neuroblastoma restraining interferon pathway activity and chemokine expression. Oncoimmunology. 2017;6(6):e1320626.

7. Mina M, Boldrini R, Citti A, Romania P, D’Alicandro V, De Ioris M, et al. Tumor-infiltrating T lymphocytes improve clinical outcome of therapy-resistant neuroblastoma. Oncoimmunology. 2015;4(9):e1019981.

8. Berthold F, Spix C, Kaatsch P, Lampert F. Incidence, Survival, and Treatment of Localized and Metastatic Neuroblastoma in Germany 1979–2015. Pediatric Drugs. 2017;19(6):577–93.

9. Simon T, Berthold F, Borkhardt A, Kremens B, De Carolis B, Hero B. Treatment and outcomes of patients with relapsed, high-risk neuroblastoma: results of German trials. Pediatr Blood Cancer. 2011;56(4):578–83.

10. Sadelain M, Rivière I, Brentjens R. Targeting tumours with genetically enhanced T lymphocytes. Nat Rev Cancer. 2003;3(1):35–45.

11. Kunkele A, Johnson AJ, Rolczynski LS, Chang CA, Hoglund V, Kelly-Spratt KS, et al. Functional Tuning of CARs Reveals Signaling Threshold above Which CD8+ CTL Antitumor Potency Is Attenuated due to Cell Fas-FasL-Dependent AICD. Cancer Immunol Res. 2015;3(4):368–79.

12. Kunkele A, Taraseviciute A, Finn LS, Johnson AJ, Berger C, Finney O, et al. Preclinical Assessment of CD171-Directed CAR T-cell Adoptive Therapy for Childhood Neuroblastoma: CE7 Epitope Target Safety and Product Manufacturing Feasibility. Clin Cancer Res. 2017;23(2):466–77.

13. Park JR, Digiusto DL, Slovak M, Wright C, Naranjo A, Wagner J, et al. Adoptive transfer of chimeric antigen receptor re-directed cytolytic T lymphocyte clones in patients with neuroblastoma. Molecular therapy : the journal of the American Society of Gene Therapy. 2007;15(4):825–33.

14. Grunewald L, Lam T, Andersch L, Klaus A, Schwiebert S, Winkler A, et al. A Reproducible Bioprinted 3D Tumor Model Serves as a Preselection Tool for CAR T Cell Therapy Optimization. Frontiers in immunology. 2021;12(2382).

15. Tjaden B, Baum K, Marquardt V, Simon M, Trajkovic-Arsic M, Kouril T, et al. N-Myc-induced metabolic rewiring creates novel therapeutic vulnerabilities in neuroblastoma. Scientific Reports. 2020;10(1):7157.

16. Andersch L, Radke J, Klaus A, Schwiebert S, Winkler A, Schumann E, et al. CD171- and GD2- specific CAR-T cells potently target retinoblastoma cells in preclinical in vitro testing. BMC Cancer. 2019;19(1):895.

17. Wang X, Naranjo A, Brown CE, Bautista C, Wong CW, Chang WC, et al. Phenotypic and functional attributes of lentivirus-modified CD19-specific human CD8+ central memory T cells manufactured at clinical scale. J Immunother. 2012;35(9):689–701.

18. Ali S, Toews K, Schwiebert S, Klaus A, Winkler A, Grunewald L, et al. Tumor-Derived Extracellular Vesicles Impair CD171-Specific CD4+ CAR T Cell Efficacy. Frontiers in immunology. 2020;11(531).

19. Lodrini M, Sprüssel A, Astrahantseff K, Tiburtius D, Konschak R, Lode HN, et al. Using droplet digital PCR to analyze MYCN and ALK copy number in plasma from patients with neuroblastoma. Oncotarget. 2017;8(49):85234–51.

20. Fuchs S, Danßmann C, Klironomos F, Winkler A, Fallmann J, Kruetzfeldt L-M, et al. Defining the landscape of circular RNAs in neuroblastoma unveils a global suppressive function of MYCN. Nature communications. 2023;14(1):3936.

21. Hartlieb SA, Sieverling L, Nadler-Holly M, Ziehm M, Toprak UH, Herrmann C, et al. Alternative lengthening of telomeres in childhood neuroblastoma from genome to proteome. Nature communications. 2021;12(1):1269.

22. Zeid R, Lawlor MA, Poon E, Reyes JM, Fulciniti M, Lopez MA, et al. Enhancer invasion shapes MYCN-dependent transcriptional amplification in neuroblastoma. Nat Genet. 2018;50(4):515–23.

23. Bosse KR, Raman P, Zhu Z, Lane M, Martinez D, Heitzeneder S, et al. Identification of GPC2 as an Oncoprotein and Candidate Immunotherapeutic Target in High-Risk Neuroblastoma. Cancer Cell. 2017;32(3):295–309.e12.

24. Landt SG, Marinov GK, Kundaje A, Kheradpour P, Pauli F, Batzoglou S, et al. ChIP-seq guidelines and practices of the ENCODE and modENCODE consortia. Genome research. 2012;22(9):1813–31.

25. Ianevski A, He L, Aittokallio T, Tang J. SynergyFinder: a web application for analyzing drug combination dose–response matrix data. Bioinformatics. 2017;33(15):2413–5.

26. Bliss CI. The toxicity of poisons applied jointly. Ann Appl Biol. 1939;26(3):585–615.

27. Zervos AS, Gyuris J, Brent R. Mxi1, a protein that specifically interacts with Max to bind Myc-Max recognition sites. Cell. 1993;72(2):223–32.

28. Zhang W, Yu Y, Hertwig F, Thierry-Mieg J, Zhang W, Thierry-Mieg D, et al. Comparison of RNA-seq and microarray-based models for clinical endpoint prediction. Genome Biology. 2015;16(1):133.

29. Popov N, Schulein C, Jaenicke LA, Eilers M. Ubiquitylation of the amino terminus of Myc by SCF(beta-TrCP) antagonizes SCF(Fbw7)-mediated turnover. Nat Cell Biol. 2010;12(10):973–81.

30. Textor A, Grunewald L, Anders K, Klaus A, Schwiebert S, Winkler A, et al. CD28 Co-Stimulus Achieves Superior CAR T Cell Effector Function against Solid Tumors Than 4-1BB Co-Stimulus. Cancers (Basel). 2021;13(5):1050.

31. Demidenko E, Miller TW. Statistical determination of synergy based on Bliss definition of drugs independence. PLoS One. 2019;14(11):e0224137.

32. Lee N, Zakka LR, Mihm MC, Jr., Schatton T. Tumour-infiltrating lymphocytes in melanoma prognosis and cancer immunotherapy. Pathology. 2016;48(2):177–87.

33. Mahmoud SM, Paish EC, Powe DG, Macmillan RD, Grainge MJ, Lee AH, et al. Tumor-infiltrating CD8+ lymphocytes predict clinical outcome in breast cancer. J Clin Oncol. 2011;29(15):1949–55.

34. Asgharzadeh S, Salo JA, Ji L, Oberthuer A, Fischer M, Berthold F, et al. Clinical significance of tumor-associated inflammatory cells in metastatic neuroblastoma. J Clin Oncol. 2012;30(28):3525–32.

35. Larsson K, Kock A, Idborg H, Arsenian Henriksson M, Martinsson T, Johnsen JI, et al. COX/mPGES-1/PGE2 pathway depicts an inflammatory-dependent high-risk neuroblastoma subset. Proceedings of the National Academy of Sciences of the United States of America. 2015;112(26):8070–5.

36. Rached J, Nasr Z, Abdallah J, Abou-Antoun T. L1-CAM knock-down radiosensitizes neuroblastoma IMR-32 cells by simultaneously decreasing MycN, but increasing PTEN protein expression. International journal of oncology. 2016;49(4):1722–30.

37. Walker AJ, Majzner RG, Zhang L, Wanhainen K, Long AH, Nguyen SM, et al. Tumor Antigen and Receptor Densities Regulate Efficacy of a Chimeric Antigen Receptor Targeting Anaplastic Lymphoma Kinase. Molecular therapy : the journal of the American Society of Gene Therapy. 2017;25(9):2189–201.

38. Ramakrishna S, Highfill SL, Walsh Z, Nguyen SM, Lei H, Shern JF, et al. Modulation of Target Antigen Density Improves CAR T-cell Functionality and Persistence. Clinical Cancer Research. 2019;25(17):5329–41.

39. Majzner RG, Rietberg SP, Sotillo E, Dong R, Vachharajani VT, Labanieh L, et al. Tuning the Antigen Density Requirement for CAR T Cell Activity. Cancer Discov. 2020.

40. Watanabe K, Terakura S, Martens AC, van Meerten T, Uchiyama S, Imai M, et al. Target antigen density governs the efficacy of anti-CD20-CD28-CD3 ζ chimeric antigen receptor-modified effector CD8+ T cells. Journal of immunology (Baltimore, Md : 1950). 2015;194(3):911-20.

41. Mossé YP, Fox E, Teachey DT, Reid JM, Safgren SL, Carol H, et al. A Phase II Study of Alisertib in Children with Recurrent/Refractory Solid Tumors or Leukemia: Children’s Oncology Group Phase I and Pilot Consortium (ADVL0921). Clinical Cancer Research. 2019;25(11):3229–38.

42. Maris JM, Morton CL, Gorlick R, Kolb EA, Lock R, Carol H, et al. Initial testing of the aurora kinase A inhibitor MLN8237 by the Pediatric Preclinical Testing Program (PPTP). Pediatr Blood Cancer. 2010;55(1):26–34.

43. Wu X, Nelson M, Basu M, Srinivasan P, Lazarski C, Zhang P, et al. MYC oncogene is associated with suppression of tumor immunity and targeting Myc induces tumor cell immunogenicity for therapeutic whole cell vaccination. Journal for ImmunoTherapy of Cancer. 2021;9(3):e001388.

