## Supplemental Text for "Targeting MYCN upregulates L1CAM tumor antigen in MYCN-dysregulated neuroblastoma to increase CAR T cell efficacy"

### Supplementary Figures

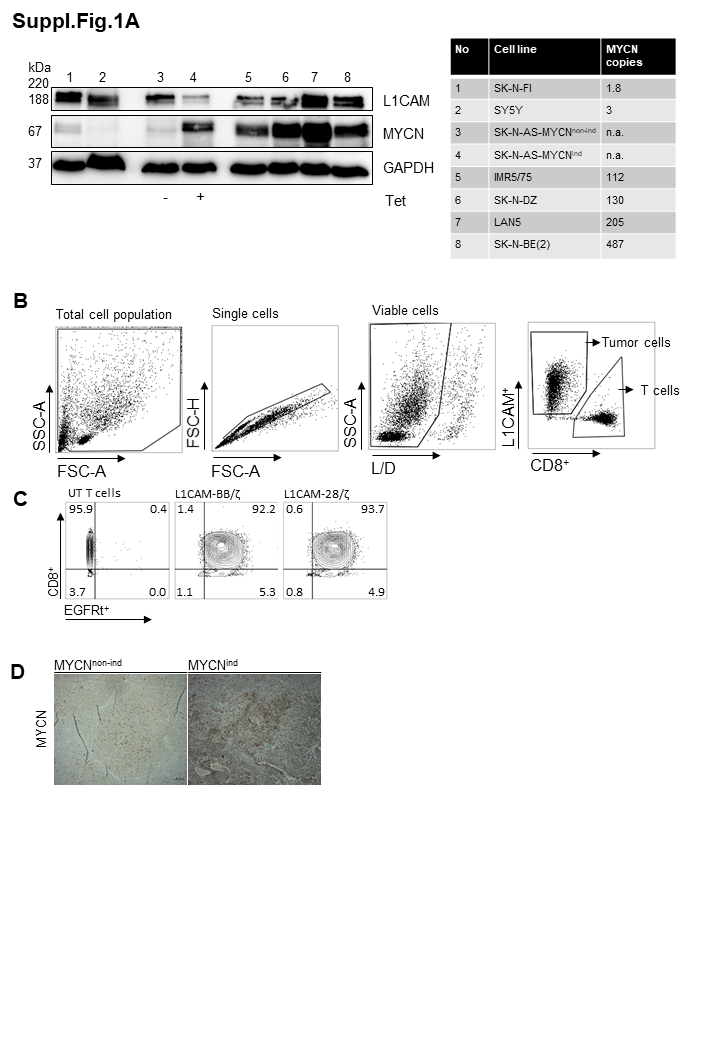

**Fig. S1: SK-N-AS-MYCN tumor cell model and gating strategy. A.** MYCN protein expression evaluated by immunoblot of neuroblastoma cell lines and tetracycline inducible tumor model with varying MYCN copy numbers: 1 SK-N-FI (1.8 copies), 2 SY5Y (3 copies), 3 SK-N-AS-MYCN^non-ind^ (n.a.), 4 SK-N-AS-MYCN^ind^ (n.a.), 5 IMR5/75 (112 copies), 6 SK-N-DZ (130 copies), 7 Lan5 (205 copies) and 8 SK-N-BE(2) (487 copies). Genomic MYCN copy numbers were published by Lodrini et al. (18). GAPDH served as loading control. **B**. Gating strategy for flow cytometry analysis throughout the experimental set-up. Gates were set as following: total cell population, single cells, viable cells and either L1CAM-positive cells for tumor cell analysis or CD8-positive cells for T cell analysis. **C.** L1CAM-specific CAR T cells were enriched for EGFRt expression as surrogate marker for CAR T cell expression, subsequently stained for EGFRt^+^ and CD8^+^ to evaluate purity of cell population by flow cytometry. Gates are set on viable, single cells. **D.** Representative immunohistochemistry staining of xenotransplanted SK-N-AS-MYCN^non-ind^ and -MYCN^ind^ samples to confirm MYCN induction. scale bar= 100µm.

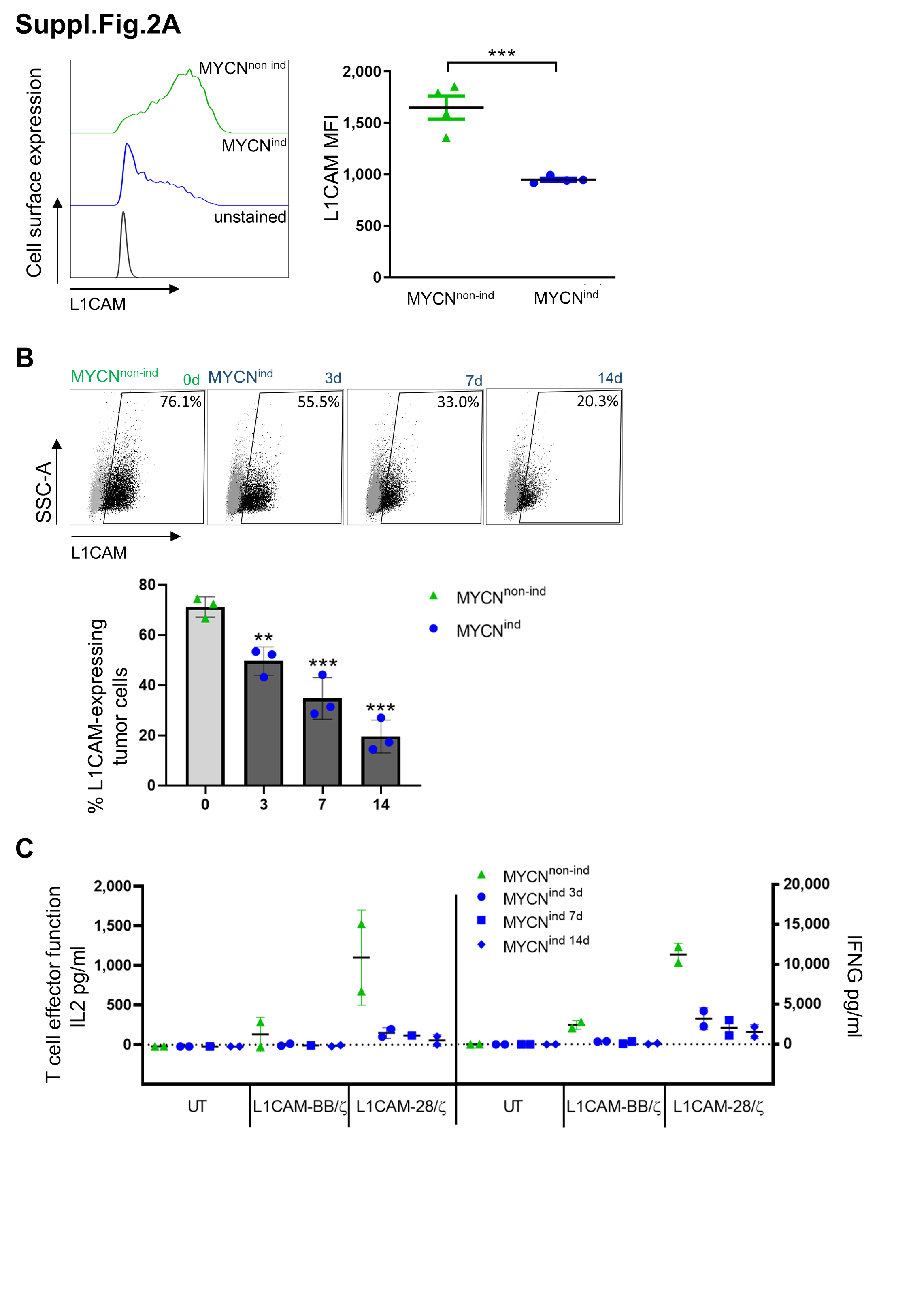

**Fig. S2: L1CAM surface expression decreases with increased time of MYCN overexpression. A.** Representative staggered histogram of flow cytometry analysis shows L1CAM expression on SK-N-AS-MYCN^non-ind/ind^ cells. MFI of L1CAM expression of individual experiments is represented as dot blots (n=3). **B.** Flow cytometry analysis of L1CAM expression on SK-N-AS-MYCN neuroblastoma cell line cultured in the presence of 2 µg/ml tetracycline for 0, 3, 7 and 14 days. Exemplary dot blots and summarized data are shown (n= 3 biological replicates, one-way ANOVA). **C.** Antigen titering by consistent treatment of SK-N-AS-MYCN^non-ind/ind^ tumor cells with tetracycline. IL2 and IFNG release by L1CAM-specific CAR T cells after 24 h-coculture with SK-N-AS-MYCN^non-ind/ind^ tumor cells (E:T 1:5, n=2 biological replicates in technical triplicates). Depicted is mean of biological triplicates/duplicates with error bars representing mean ± SD, *, p≤0.5; **, p≤0.01, ***, p≤0.001

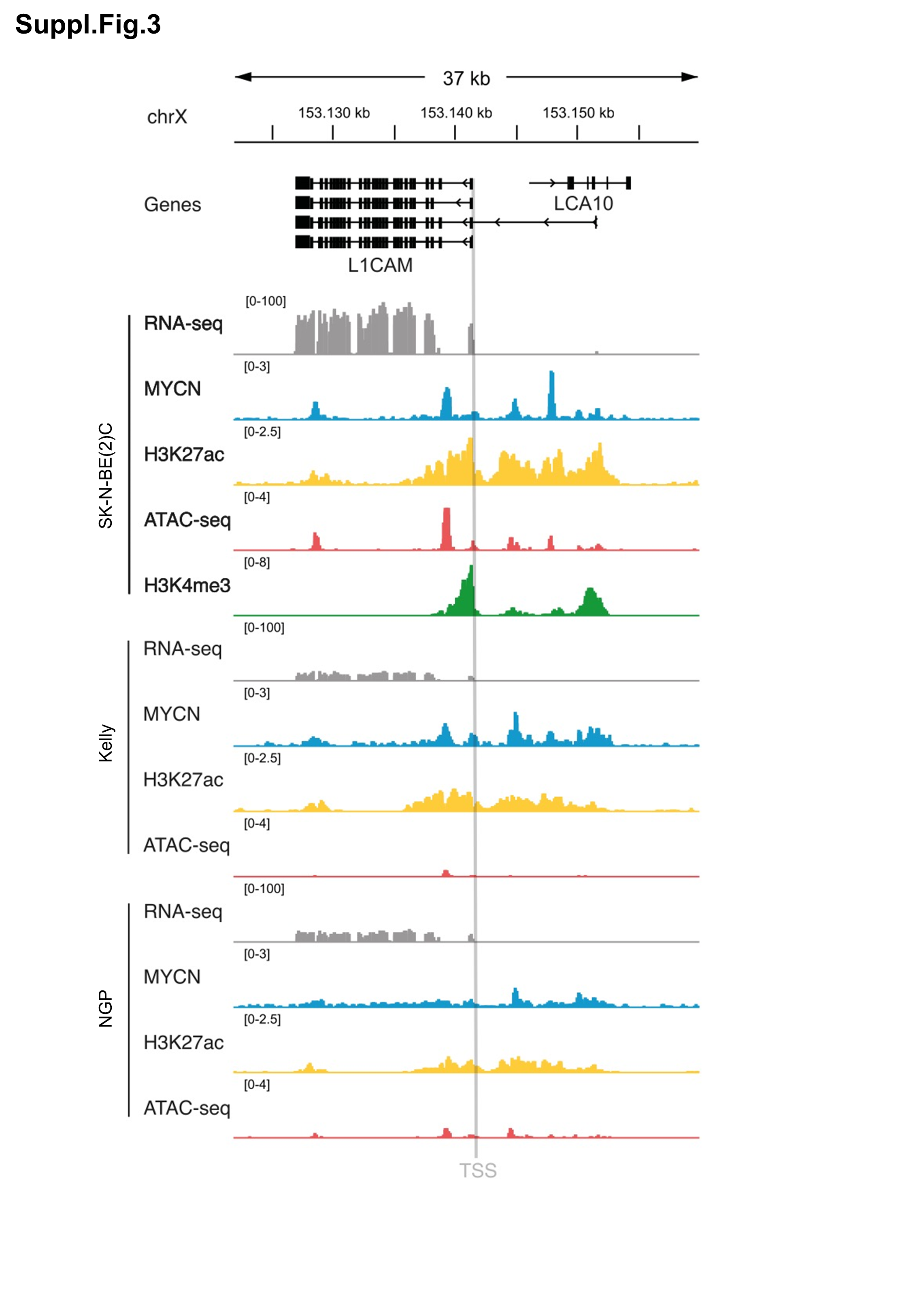

**Fig. S3: ChIP-sequencing data reveals binding of MYCN to L1CAM locus.** Relative occurrence of MYCN ChIP-seq profiles on *L1CAM* close to transcriptional start site (TSS, grey) for three *MYCN*-amplified neuroblastoma cell lines NGP, Kelly and SK-N-BE(2)-C.

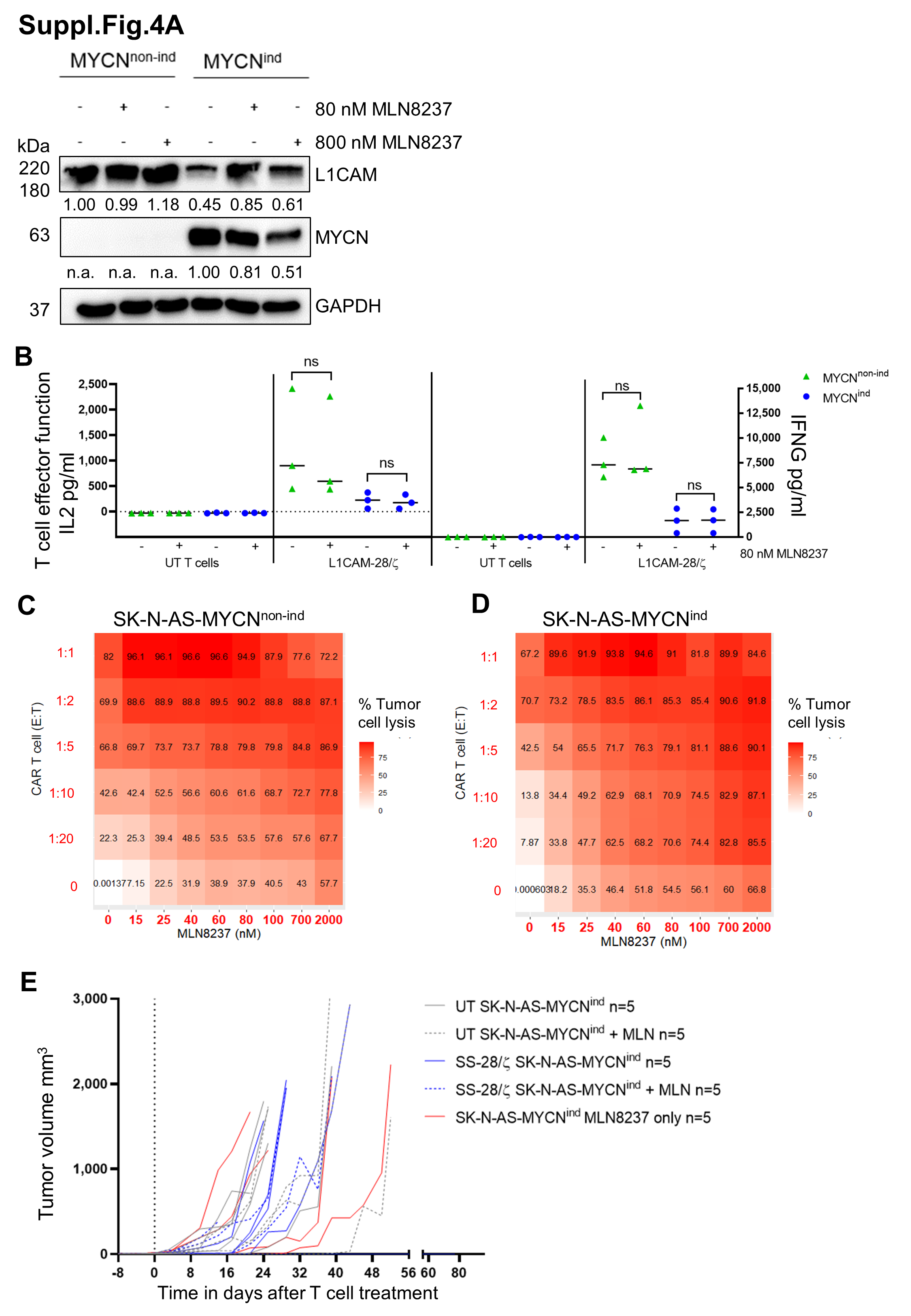

**Fig. S4: Synergism of L1CAM-CAR T cells and MLN8237 in SK-N-AS-MYCN^ind^ tumor cells. A.** Immunoblot evaluating MYCN and L1CAM expression of SK-N-AS-MYCN^non-ind^ and -MYCN^ind^ tumor cells treated with 80 or 800nM MLN8237 over 72 hours. GAPDH serves as loading control. **B.** IL2 and IFNG release by L1CAM-specific CAR T cells cocultured with SK-N-AS-MYCN^non-ind^ and -MYCN^ind^ tumor cells in addition with 80nM MLN8237 (E:T 1:5, 24 h). Depicted is mean of biological triplicates with error bars representing mean ± SD, ns = not significant **C.** and **D.** Heatmaps indicating tumor cell lysis for each dose-combination against **C.** SK-N-AS-MYCN^non-ind^ and **D.** SK-N-AS-MYCN^ind^ tumor cells, calculated with the Bliss synergy score using R. **E.** Tumor growth of SK-N-AS-MYCN^ind^ tumors in NSG mice treated with untransduced (UT) and L1CAM-28/ζ CAR T cells alone or in combination with MLN8237 (all groups n=5). Zero (“0”) indicates start of treatment.
